# Logical model of human tolerogenic dendritic cells and their participation in autoimmune disease

**DOI:** 10.1101/2023.08.22.554293

**Authors:** Karen J. Nuñez-Reza, Isaac Lozano-Jiménez, Leslie Martínez-Hernández, Alejandra Medina-Rivera

## Abstract

Tolerogenic dendritic cells (tolDC) regulate the immune response, several clinical trials focused on autoimmune diseases use tolDC to promote immune tolerance response and Treg activation. Here we built a logical model for the tolerization cellular process of dendritic cells using IL10. By combining literature knowledge, microarray gene expression, and key tolDC markers, we ensembled a logical model that describes the obtention of tolDC using the IL10 signaling cascade that spawns the most tolerogenic phenotype. The model uses IL10 as input and the signaling cascade that trigger seven transcription factors (TFs), three previously known TFs in the IL10 response (STAT3, NFKB, STAT6), and four were incorporated based on our gene expression analysis (IRF8, TCF7L2, CEBPB, and TFCP2L1). Using our model, we generated mutants *in-silico* and identified that even when IL10 is present the single mutants for TCF7L2, IRF8, TFCP2L1, and STAT3 were not able to reach a tolDC stable state, highlighting the relevance of these TFs in the process. The current model sets a precedent that will help in the development of tolDC for future applications.

## Introduction

Tolerogenic dendritic cells (tolDC) are part of the system that modulates the immune response and are characterized by the decreased expression of co-stimulatory molecules, the upregulation of inhibitory molecules, and the augmented production of anti-inflammatory cytokines (1). TolDC often displays an immature or semi-mature phenotype that is characterized by altered cytokine production (2). The presentation of low levels of antigen without co-stimulation leads to T cell anergy and promotion of regulatory T cell differentiation (3).

The current treatments for autoimmune diseases are based on immune suppressors and have several side effects. TolDC are being tested as antigen-specific immunotherapy for several autoimmune diseases, due to their ability to promote immune tolerance response and to prime Treg (1). Li et al. (4) demonstrated the development of a novel approach to combine tolDC and mesenchymal stem cells as immunotherapy for arthritis. Odobasic et al. (5) generated tolDC to suppress glomerulonephritis caused by autoimmunity against myeloperoxidase. Immune tolerance has emerged as an alternative for the future development of treatments for autoimmune and allergic diseases (3).

There are several methods to obtain tolDC *in vitro*, that used dexamethasone, IL10, rapamycin, vitamin D, and/or TNFα. However, IL10 differentiation exhibits the most robust tolerogenic phenotype by inducing Treg which suppressed T cell reactivity (6,7).

Advances in mathematics, biology, and computer science have led to the development of various techniques that allow *in-silico* modeling of biological systems (8,9). Logical models are used to study many components of systems biology, including metabolism, signaling pathways, and transcription networks (10). Logical models have been used as prior knowledge for the generation of hypotheses and predictions, allowing us to make diagrams and illustrate the components and interactions of these systems to help us understand and represent the general structures of biological systems.

Our study aimed to build a logical model using healthy tolDC and identify key players in the tolerization cellular process. We built our model from literature, integrated it with microarray gene expression data, and predicted regulatory interactions from highly expressed transcription factors (TFs).

## Results

### Novel overexpressed TFs could be key in the tolDC commitment process

We searched for available public transcriptome data from tolDC obtained from moDC using IL10, to identify differentially expressed genes and look for relevant genes that could be included in the model. We found eight samples analyzed by Agilent microarrays and publicly available from GEO (GSE117946) (13). The RNA was obtained from healthy human peripheral blood CD14+ monocyte cells differentiated into moDC (IL4 and GM-CSF used) and tolDC (IL10, IL4, and GM-CSF used) (13).

We compared the transcriptome of moDC vs tolDC (see methods), and we were able to identify new actors relevant to tolDC obtention using IL10 (see Supplementary Table 2). Like the downregulation of CD1E, a gene related to antigen-presenting functions. CD163-specific marker for monocyte lineage was overexpressed in tolDC, which has been previously reported (20). We identified IL7 between the overexpressed genes in tolDC, which has a regulatory role on B cells (21). CD83 a specific marker for moDC maturation was underexpressed, confirming tolDC behavior, CD83 has been identified as overexpressed in patients with autoimmune diseases (22). Surprisingly we identified that CD52 was underexpressed in tolDC, something that is not observed in tolDC obtained with dexamethasone and vitamin D, which is a specific high-marker for IL10 differentiated tolDC () (23,24), this difference could be explained by the tolerogenic protocol used. In a study of tolDC generated by vitamin D, they identified CSF1 as a regulator of the process (25), in contrast, in our analysis we found the CSF1 gene underexpressed in tolDC.

As little is known regarding the transcriptional regulatory mechanisms acting in tolDC obtention, we set out to identify additional relevant TFs using differentially expressed gene data. We identified four overexpressed TFs in tolDC, TCF7L2, IRF8, CEBPB, and TFCP2L1 that could be involved in tolDC commitment, none of these TFs have been described in the context of tolDC commitment before. IRF8 and CEBPB have been reported to be interacting and negatively regulating the differentiation of monocytes to moDCs (26). We found KLF5 and KLF8 underexpressed in tolDC, KLF5 has been related to granulocyte differentiation (27), and KLF8 has been related to adipocyte differentiation (28). We also identified that IRF4, BATF3, CEBPA, and NR4A3 which are related to moDC differentiation (29) are underexpressed in tolDC.

### Identification of novel TF binding sites for tolDC-related genes

In order to identify the regulatory targets of the overexpressed TFs we used the position-weight matrices of the four overexpress TF (TCF7L2, IRF8, CEBPB, TFCP2L1), and the TFs that participated in IL10 signaling, IRF4, STAT3, and STAT6 and used the pattern-matching tool matrix-scan from the RSAT suite (17) to identify transcription factor binding sites (TFBS). We looked for predicted TFBS in the 2kb upstream of the transcription start site (TSS) for the genes relevant to tolDC behavior previously described (30) (Table 1).

**Table 1.** Genes that are signature of tolDC (30,31).

| **Gene** | **Function** | **tolDC** |
| --- | --- | --- |
| IL1B | Proinflammatory cytokines | Down |
| IL6 | Proinflammatory cytokines | Down |
| IL12A | Proinflammatory cytokines | Down |
| TNF | Proinflammatory cytokines | Down |
| CD80 | DC maturation | Down |
| CD86 | DC maturation | Down |
| CD40 | DC maturation | Down |
| HLA-DRA | DC maturation | Down |
| IL10 | Anti-inflammatory | Up |
| IL1RN | Anti-inflammatory | Up |
| C1QA | Anti-inflammatory | Up |
| THBS1 | Anti-inflammatory | Up |
| CCR5 | Chemokines | Up |
| CCR7 | Chemokines | Up |
| CXCR4 | Chemokines | Up |
| CCL18 | Chemokines | Up |
| CCL22 | Chemokines | Up |
| CXCL1 | Chemokines | Up |
| IDO1 | Metabolic control | Up |
| HMOX1 | Metabolic control | Up |
| ARG1 | Metabolic control | Up |
| ALDH2 | Metabolic control | Up |
| ICOSLG | Immunomodulatory molecules | Up |
| BTLA | Immunomodulatory molecules | Up |
| CTLA4 | Immunomodulatory molecules | Up |
| PDCD1 | Immunomodulatory molecules | Up |
| FASLG | Immunomodulatory molecules | Up |

We were able to identify TCF7L2 predicted TFBSs in the upstream region of the proinflammatory genes, DC maturation markers, chemokines, and anti-inflammatory genes (Table1 and Figure 2), TCF7L2 might be possibly negatively regulating moDC markers and positively regulating tolDC genes. TFCP2L1 is probably positively regulating chemokines. IRF8 is the TF with the lesser number of predicted binding sites exclusive to five genes, immunomodulators CTLA4 and HMOX1, chemokines CCL18 and CXCL1, and proinflammatory TNF. CEBPB which has been related to an inflammatory response (32) has TFBS for proinflammatory genes (IL1B, IL6, CD80, CD86, HLA-DRA). In our analysis, the number of TFs possibly regulating each gene goes from one to three. We also wanted to corroborate the predicted sites found with sites from ChIP-seq experiments from ReMap, we found ChIP-seq data for CEBPB, IRF8, and TCF7L2. The predicted sites that we were able to corroborate with ChIP-seq data are those with an asterisk in Figure 2.

**Figure 1.**
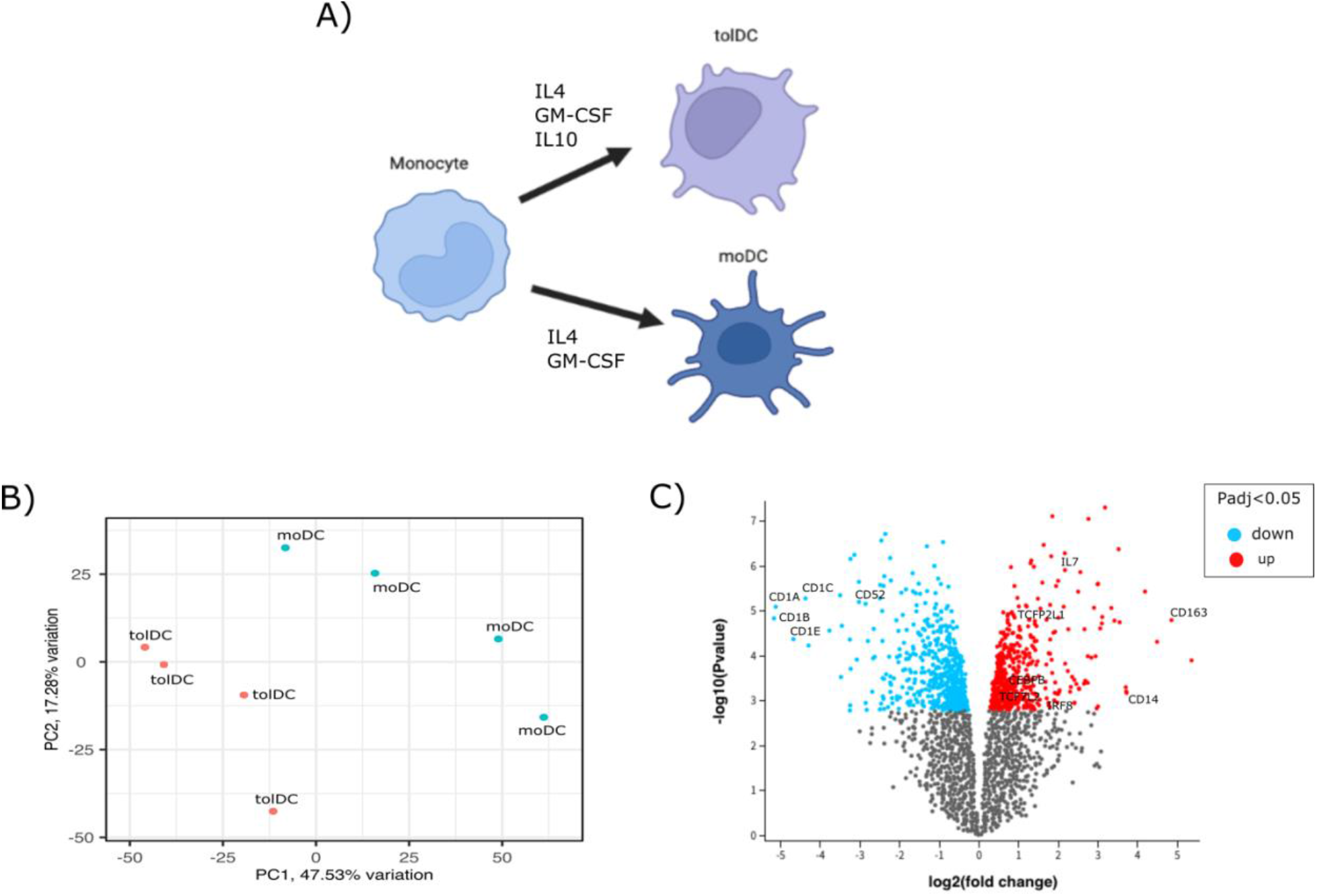
tolDC differentiation and differential expression between tolDC and moDC. A) Diagram of the monocyte differentiation to tolDC and moDC and their specific cultured cytokines. B) PCA plot of the samples used in our analysis, we can observe two separated groups. C) Volcano plot of the genes overexpressed and underexpressed into tolDC of four biological replicates.

**Figure 2.**
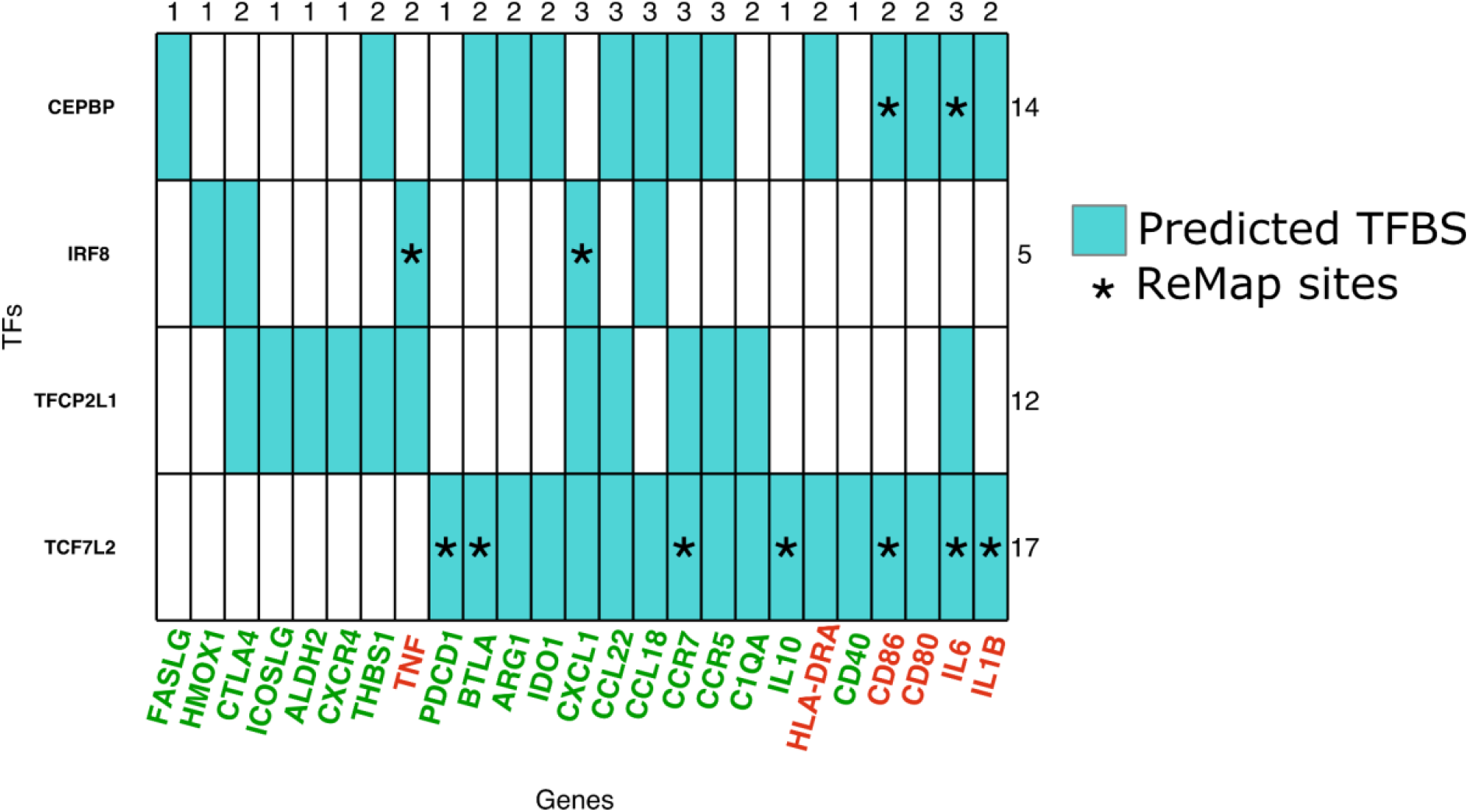
Heatmap of the TFBS predicted with our analysis. On the x-axis, genes related to inflammatory response in red and genes related to tolerogenic behavior in green, and on the y-axis four overexpressed TFs. Green squares indicate predicted TFBS and the asterisk marks interactions that were corroborated by ChIP-seq experiments from Remap database.

### The logical model of moDC to tolDC differentiation describes two stable states corresponding to each cell type

Considering the literature (Supplementary Table 3), specific moDC and tolDC markers (Table 1), analyzed microarray gene expression (Supplementary Table 2), and newly identified TF regulatory interactions (Figure 2), we built a logical model that describes the obtention of tolDC using IL10 by incorporating the IL10 signaling cascade (Figure 3).

**Figure 3.**
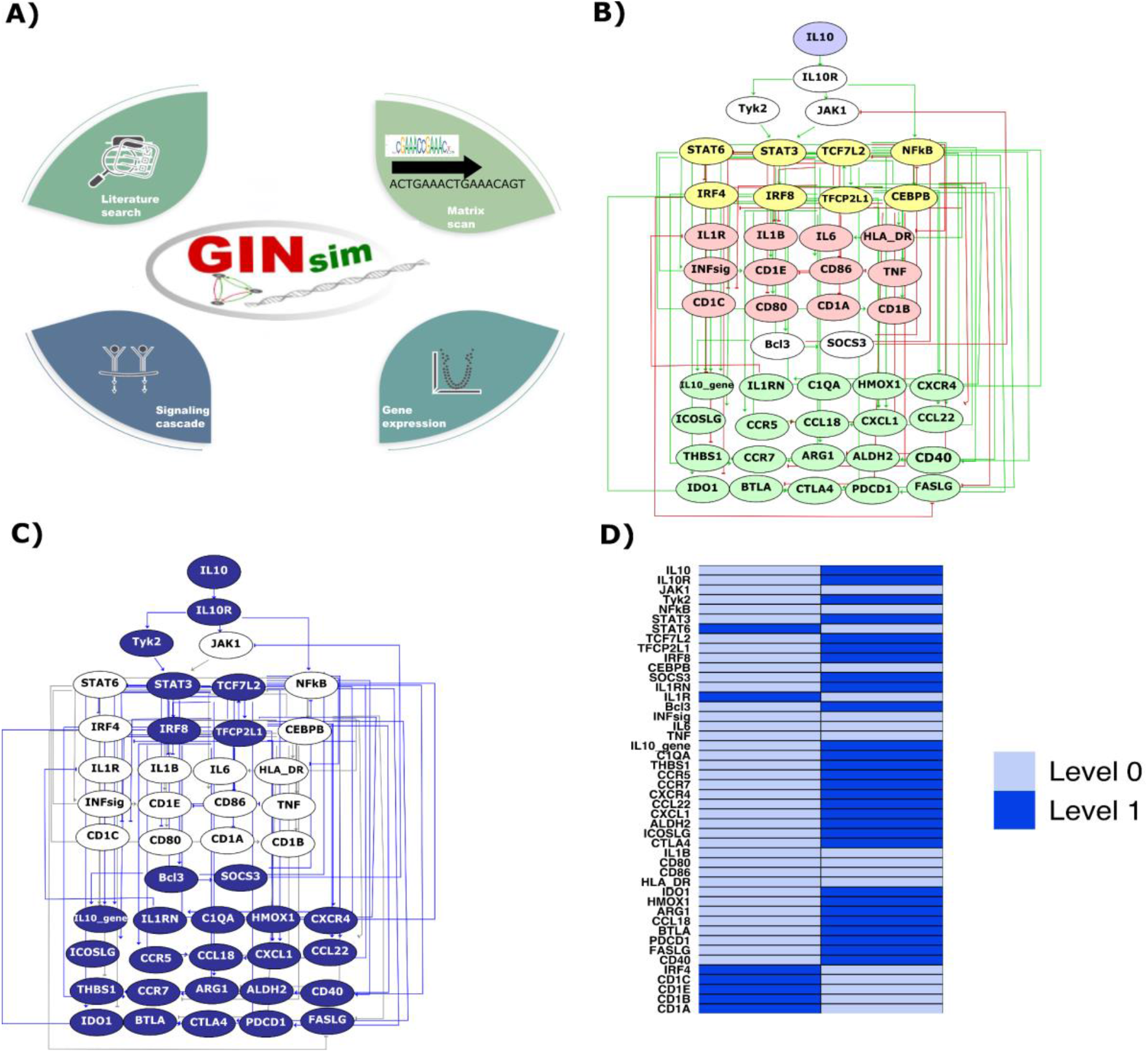
Model build of the tolerogenic dendritic cell. A) To build our model we considered the literature, predicted TFBS, gene-expression data, and the canonical IL10 signaling cascade. B) Our complete model has one input (IL10), the signaling cascade, the TFs that participate, and the final nodes that describe an immature moDC state or tolerogenic state. Nodes in white represent our input and proteins of the signaling cascade. The nodes in yellow represent TFs. Nodes in green highlight genes related to tolerogenic outcomes. Nodes in red are genes related to inflammation. C) Nodes in blue highlight genes in the stable state related to tolDC commitment.D) Stable states reached with our model, representing the final condition of every component in each state. Light blue indicates the node is OFF and dark blue indicates the node is ON.

Our logical model is composed of arcs and nodes, each node corresponds to a specific component that could be a cytokine, receptor, protein, or gene; the arcs describe the relations between the nodes, which could be negative or positive interactions. The model is built using IL10 as input, the signaling cascades that it triggers (12), the differentially expressed genes, and predicted TFBS (Figure 3A). STAT3 and NFKB expression is directly triggered by IL10, and STAT6 is negatively regulated by STAT3 which is relevant as STAT6 and STAT3 are required for moDC differentiation (29). The rest of the TFs were added as result of our transcriptome analysis (Supplementary Table 2). The nodes in red correspond to genes related to inflammation, and the nodes in green correspond to genes related to tolDC commitment. Nodes in green and red correspond to the final nodes (Figure 3B).

Once we completed the regulatory graph (Figure 3B) and set the Boolean rules describing the interaction between nodes (Supplementary Table 1), we computed the stable states. The first state with IL10 ON corresponds to tolDC, this state is characterized by inflammatory genes being OFF, and the specific genes for tolerogenic behavior having an ON setting (Figure 3C). The second state marks the absence of IL10 and corresponds to moDC commitment, characterized by the moDC-specific genes being ON and the genes specific for tolDC being OFF. In summary, the two stable states correspond to the expected outcomes for moDC and tolDC cell types of commitment (Figure 3D).

### In silico mutations reveal the importance of STAT3, TCF7L2, and TFCP2L1 for tolDC commitment

To test our model, we used the CoLoMoTo suite which gathers several software tools like Pint, BioLQM, and MaBoSS. Our analysis has been released in a Jupyter notebook available at https://github.com/karenunez/logical_model_tolDC to ensure reproducibility. We used BioLQM to search for trap spaces that correspond to cyclic attractors which indicate a partial state that is not possible to evade and could impact cellular commitment (33), in our model, there are no trap spaces.

We also used BioLQM to perform *in silico* mutations to predict their impact on the tolerization process. We mutated the TFs, STAT6, STAT3, IRF8, TCF7L2, CEBPB, TFCP2L1, NFkB, and the IL10 receptor (IL10R gene) (Figure 4A), with each perturbation stable states are calculated. When IL10 is not present (OFF) the CEBPB perturbation enables the activation of moDC-specific genes (state 0). The individual perturbations of TCF7L2, TFCP2L1, and IL10R, have the same behavior activating moDC-specific genes and repressing tolerogenic genes in the absence of IL10. The single perturbation of IL10R in the presence of IL10 turns off tolerogenic genes. STAT3 perturbation in the presence of IL10 abolishes the activation of tolerogenic genes and allows the activation of inflammatory genes. STAT3 has been reported to have an important role in obtaining tolDC with vitamin D protocol (34). TFCP2L1 perturbation in the presence of IL10 repressed the expression of several tolerogenic genes. TCF7L2 perturbation in the presence of IL10 suppressed half of the tolerogenic genes. IRF8 perturbation in the presence of IL10 suppressed the activation of the complete set of tolerogenic behavior.

**Figure 4.**
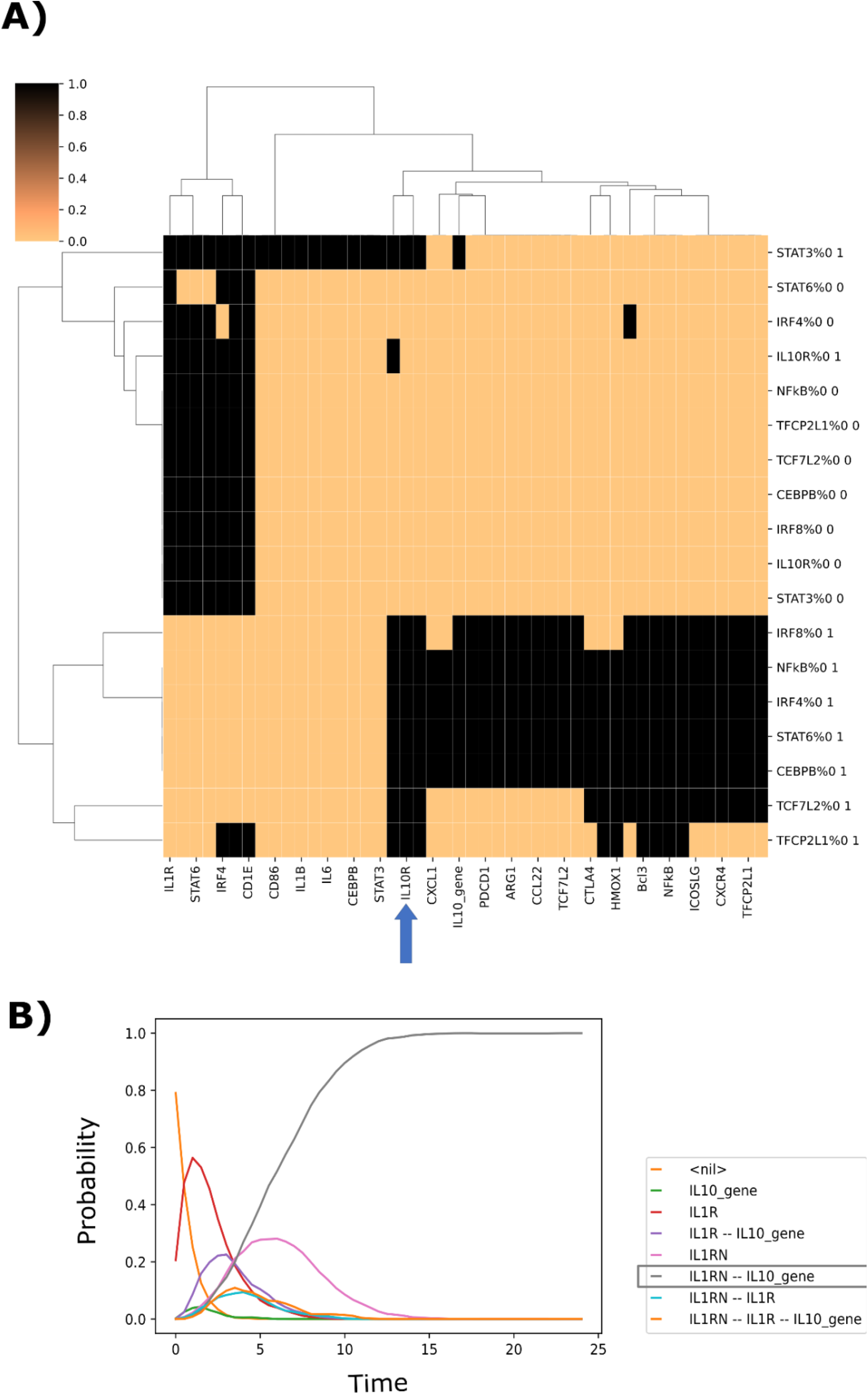
CoLoMoTo allows us to perform several mutations at a time. A) The columns represent every node, and the rows represent the stable state obtained with the mutant described, for example, “CEBPB%0 0” means knock out in CEBPB, and the state 0 without IL10, IL10R is marked with an arrow. B) Trajectories calculated with MaBoSS, the curve in grey correspond to tolDC commitment.

We wanted to know the reachability of our stable states, for this, we used the tool MaBoSS which performs Stochastic Boolean simulations by generating mean temporal curves (19). The initial conditions correspond to monocyte, the stable states are searched in the presence or absence of IL10. In Figure 4B, we examine a graph with several initial curves that represent transient cellular states from the initial state (monocyte) to the tolerogenic state (tolDC). One single curve reached level 1, which corresponds to 100% of the cell population reaching the tolDC state.

## Discussion

TolDC are important in the immune response and are becoming relevant due to their possible usage as therapy in immune diseases (35–37). Logical models have been widely used to describe biological processes like T-cell commitment (38), cell death (39), cell interaction during infection (40), sex determination (41), etc. Our model describes the tolerization of monocytes and the principal participants of this commitment.

There are several protocols to obtain tolDC, our model is focused on the protocol using IL10. Based on our gene expression analysis performed using the IL10 differentiation protocol, we identified four overexpressed TFs, IRF8, TCF7L2, CEBPB, and TFCP2L1. We predicted several TFBS that require further experimental confirmation and allowed us to add the specific tolDC genes into our model. With the integration of predicted TFBS, gene expression, and IL10 signaling cascade we were able to complete the model of dendritic cell tolerization, showing two clear final states (moDC and tolDC) and no trap spaces (intermediated differentiation states).

We identified differences between the canonical tolDC genes obtained with vitamin D3 tolerization, like CD52, that we found under expressed; the main difference being the tolerogenic protocol.

Recently a genetic variant rs2280381 has been identified in a monocyte-specific enhancer. The SNP rs2280381 allows the overexpression of the IRF8 gene in monocytes, whereas patients with SLE have a risk allele T that underexpressed the IRF8 gene (42). This could impact the differentiation of tolDC, as we identify it as essential for tolDC differentiation.

Our model was tested under different mutations and predicted the role of STAT3, IRF8, STAT6, IRF4, NFkB, CEBPB, TCF7L2, and TFCP2L1 in tolDC commitment under IL10 differentiation. We also corroborated the role of STAT3 as the major regulator, we identified that a perturbation on the TF allows the activation of inflammatory genes. IRF8, TCF7L2, and TFCP2L1 are also participating in the tolerization process, by activation of the tolerogenic genes in addition to STAT3. According to our analysis, we identified that IRF8, STAT3, TCF7L2, and TFCP2L1 are the key regulators for the tolerization process triggered by IL10 stimulation. Our model set the foundation for experimental corroboration and integrating the different tolerization protocols. Our model can be used to evaluate the efficacy of gene therapy in the treatment of autoimmune diseases.

## Materials and Methods

### Logical model implementation

In general, logical models are composed of nodes and arcs. The nodes accept values of zero or one and could represent a protein, gene, or TF, additionally, the arcs represent interaction between the nodes, those interaction can be positive or negative. The model is set through logical rules or functions, those logical rules were stablished using Boolean operators AND (&), OR (|), and NOT (!) (11). We used GINsim 3.0 (11) software to build the regulatory network. We used previously described IL10 signaling cascade in cultured monocytes from healthy donors as a scaffold to build our model from the study by Schülke S (12). Additionally, we reviewed the tolDC differentiation literature in PubMed database, we used the terms “IL10” and “tolDC”. The data reviewed was utilized to define the logical rules of interactions between the nodes (Supplementary Table 1). We also define the rules considering the gene expression (see below). To set the TF effect, we searched the literature to determine if the regulation could be negative or positive (using the terms “TF” and “tolDC”). The computation of stable states was performed using GINsim (11).

### Pipeline for microarray analysis

We searched in the Gene Expression Omnibus (GEO) repository for available data for tolDC obtention solely with IL10 (we used the terms “tolDC” and “IL10”), and we found one series: GSE117946 which is an Affymetrix Human Gene 1.0 ST Array (13). We followed the GEO R pipeline to download the dataset (14). In brief, we used the GEOquery R package version 2.58.0 to import GEO data into the R environment. To identify differentially expressed genes between monocyte-derived DC (moDC) and tolDC, we used the limma package (version 3.46.0) to calculate log2 transformation, set the contrast with makeContrasts(), and computed statistics with Ebayes() function to identify the top differentially expressed genes and we calculated precision weights to adjust the variance. To calculate the adjusted P-value we used Benjamini & Hochberg method (p-adj 0.01) (15). The code is available on GitHub (https://github.com/karenunez/logical_model_tolDC).

Recently, a tolDC signature was published based on tolDC gene expression data (upregulated genes: CEBPB, SMPDL3A, NOD2, CD14, PAPSS2, ST3GAL1, SEMA6B, CD300LF, ACSL1, TREM1, CD93, NCOA4, C1QA, and downregulated genes: IRF4, TRIM36, MTCL1, HCAR2, MMP12, KCTD6, ZFP69, PP1R16A, CD1A, CD1E, CD1B, CD1C, IL1RAP), where authors gather all the studies available regardless of the protocol used to obtain tolDC (16). According to their data, we identified the same differentially expressed genes in dataset GSE117946 corresponding to tolDC IL10 differentiation.

### Identification of transcription factor binding sites

In order to obtain the upstream region of the selected genes, we used the tool retrieve-ensembl-seq from the RSAT suite (17), with the following options, specie: Homo sapiens, Ensembl database version 106, single organism, list of genes: IL1B, IL6, IL12A, TNF, CD80, CD86, CD40, HLA-DRA, IL10, IL11RN, C1QA, THBS1, CCR5, CCR7, CXCR4, CCL18, CCL22, CXCL1, IDO1, HMOX1, ARG1, ALDH2, ICOSLG, BTLA, CTLA4, PDCD1, CD1A, CD1B, CD1C, CD1E, IRF4, and FASLG, sequence position: upstream, from -2000 to -1, relative feature: gene. To obtain the position weight matrices, we used the tool retrieve-matrix using Jaspar version 2022 and selected the transfac format. To perform pattern matching with the matrices for four differentially expressed TFs (IRF8, TCF7L2, CEBPB, TFCP2L1, IRF4, STAT6, STAT3) to connect them to the model, we used the matrix-scan-quick tool, also available in the RSAT suite, with the following parameters - pseudo 1 -decimals 1 -2str -origin end -bginput -matrix_format transfac -markov 2 -bg_pseudo 0.01 -return sites -lth score 1, p-value 1e-4. To corroborate our discovered sites, we took all the sites found in ChIP-seq data from the Remap database and found only experiments for tree TFs. The matrices and TSS regions used are available on GitHub (https://github.com/karenunez/logical_model_tolDC).

### Notebook implementation

With the aim of achieve reproducibility, we used the CoLoMoTo platform version colomoto-docker:2022-07-01 (18) which gathers several logical modeling tools including GINsim, bioLQM, Pint, and MaBoSS. GINsim allowed us to compute stable states, and bioLQM helped us to search for trap-spaces corresponding to cyclic attractors. MaBoSS computed mean stochastic temporal trajectories of the cellular commitment (19). The GINsim model and the CoLoMoTo notebook are available in a GitHub repository (https://github.com/karenunez/logical_model_tolDC). The CoLoMoTo notebook results can be reproduced within the CoLoMoTo Docker image following the instructions provided at http://colomoto.org/notebook; in brief, go to mybinder.org/v2/gh/colomoto/colomoto-docker/mybinder/latest, then upload the notebook (tolorezation_IL10.ipynb) and the model (model_tolerization_IL10.zginml) in mybinder, then start the notebook and run it.

## Acknowledgments

This work received technical support from Luis Aguilar, Alejandro León, and Jair García of the Laboratorio Nacional de Visualización Científica Avanzada. We thank Carina Uribe Díaz, and Alejandra Castillo Carbajal for their technical support. We thank Ana Beatriz Villaseñor-Altamirano and Celine Hernandez for their comments and review of this work.

## Author contributions

ILJ, LMH, AMR, and KNR led the study and wrote the manuscript. ILJ, LMH, and KNR analyzed the data. All authors contributed to the article design and drafting of the manuscript.

## Data availability

The datasets and computer code produced in this study are available in the following databases:

- Modeling computer scripts: GitHub (https://github.com/karenunez/logical_model_tolDC)

