## Supplementary Table 1 for "Logical model of human tolerogenic dendritic cells and their participation in autoimmune disease"

Supplementary Table 1. Logical rules of the Model of Tolerization of Monocytes using IL10

| **Component** | **Logical Rule** |
| --- | --- |
| ALDH2 | TFCP2L1&GAL4 |
| ARG1 | TCF7L2&!CEBPB |
| BTLA | TCF7L2&!CEBPB |
| Bcl3 | STAT3 |
| C1QA | TCF7L2&TFCP2L1&GAL4 |
| CCL18 | !IRF8 |
| CCL22 | TCF7L2&TFCP2L1&!CEBPB |
| CCR5 | TFCP2L1&!CEBPB |
| CCR7 | TFCP2L1&!CEBPB |
| CD1A | !TFCP2L1&!GAL4 |
| CD1B | IRF8&CEBPB&!GAL4 |
| CD1C | !TCF7L2&!TFCP2L1 |
| CD1E | !TFCP2L1&CEBPB |
| CD40 | TCF7L2&GAL4 |
| CD80 | !TCF7L2&CEBPB |
| CD86 | !TCF7L2&CEBPB |
| CEBPB | NFkB |
| CTLA4 | TFCP2L1&!IRF8&GAL4 |
| CXCL1 | TCF7L2&TFCP2L1&!IRF8 |
| CXCR4 | GAL4 |
| ERK | !IL10&!SOCS3 |
| FASLG | !CEBPB |
| GAL4 | TFCP2L1&!CEBPB |
| HLA_DR | !TCF7L2&CEBPB&!GAL4 |
| HMOX1 | !IRF8&GAL4 |
| ICOSLG | TFCP2L1&GAL4 |
| IDO1 | TCF7L2&!CEBPB |
| IL10 | IL10 |
| IL10R | IL10 |
| IL10_gene | !STAT6&NFkBp50p50&Bcl3 |
| IL12 | STAT6&!TCF7L2&!TFCP2L1&!GAL4 |
| IL1B | !TCF7L2&!TFCP2L1&CEBPB&!GAL4 |
| IL1R | !IL1RN |
| IL1RN | STAT3 |
| IL6 | !TCF7L2&!TFCP2L1&CEBPB&!GAL4 |
| INFsig | NFkB |
| IRF4 | !TCF7L2&!TFCP2L1&!GAL4 |
| IRF8 | ERK |
| JAK1 | IL10R&!SOCS3 |
| NFkB | IL1R&!Bcl3 |
| NFkBp50p50 | IL10 |
| PDCD1 | TCF7L2 |
| SOCS3 | STAT3 |
| STAT3 | !JAK1&Tyk2 \| JAK1 |
| STAT6 | !STAT3 |
| TCF7L2 | TFCP2L1&!IRF8&!CEBPB |
| TFCP2L1 | STAT3 |
| THBS1 | !TFCP2L1&CEBPB&!GAL4 |
| TNF | !TFCP2L1&IRF8&CEBPB |
| Tyk2 | IL10R |

*&: AND*

*!: NOT*

*|: OR*
