## Supplementary Table 3 for "Logical model of human tolerogenic dendritic cells and their participation in autoimmune disease"

Supplementary table 3. Documentation of the model "Tolerization_model_IL10"

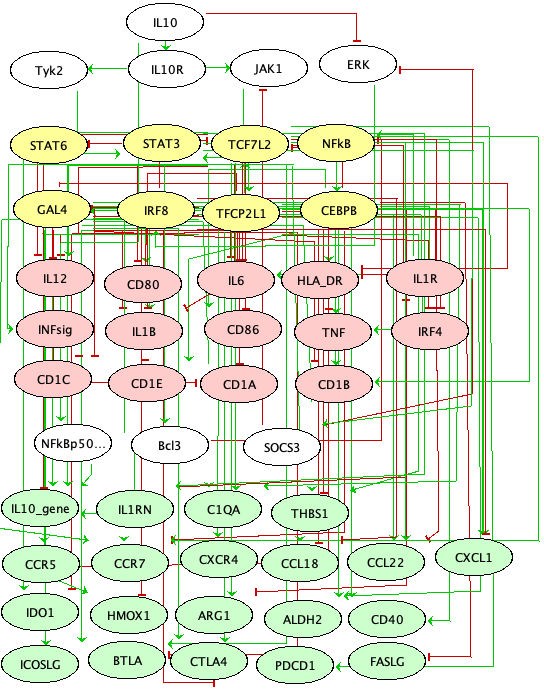

| **Nodes ID** | **Val** | **Logical function** |
| --- | --- | --- |
| IL10 | Input node | |
| IL-10 homodimers bind to the extracellular portion of the IL-10R receptor.  Schülke S. Induction of Interleukin-10 Producing Dendritic Cells As a Tool to Suppress Allergen-Specific T Helper 2 Responses. Front Immunol. 2018 Mar 19;9:455. doi: 10.3389/fimmu.2018.00455. PMID: 29616018; PMCID: PMC5867300. | |  |
| IL10R | 1 | - IL10 |
| After the union of IL-10 homodimers, the recruitment of Jak1 and its subsequent phosphorylation by the alpha chain is triggered, while Tyk2 is recruited and similarly phosphorylated by the beta chain.   1. Schülke S. Induction of Interleukin-10 Producing Dendritic Cells As a Tool to Suppress Allergen-Specific T Helper 2 Responses. Front Immunol. 2018 Mar 19;9:455. doi: 10.3389/fimmu.2018.00455. PMID: 29616018; PMCID: PMC5867300. | |  |
| JAK1 | 1 | - IL10R & !SOCS3 |
| The phosphorylation of the Jak1-Tyk2 complex will mediate the phosphorylation of STAT3.   1. Schülke S. Induction of Interleukin-10 Producing Dendritic Cells As a Tool to Suppress Allergen-Specific T Helper 2 Responses. Front Immunol. 2018 Mar 19;9:455. doi: 10.3389/fimmu.2018.00455. PMID: 29616018; PMCID: PMC5867300. | |  |
| Tyk2 | 1 | - IL10R |
| Phosphorylation of the Jak1/Tyk2 complex will mediate STAT3 phosphorylation..   1. Schülke S. Induction of Interleukin-10 Producing Dendritic Cells As a Tool to Suppress Allergen-Specific T Helper 2 Responses. Front Immunol. 2018 Mar 19;9:455. doi: 10.3389/fimmu.2018.00455. PMID: 29616018; PMCID: PMC5867300. | |  |
| STAT3 | 1 | - Tyk2 \| JAK1 |
| Once STAT3 is activated and phosphorylated, it causes a translocation of its homodimers to the nucleus where they bind to STAT binding elements and induce the expression of genes that respond to STAT3. These include suppressor of cytokine signaling 3 (SOCS-3), IL-1 receptor antagonist, transcription factors such as Bcl3.  It also inhibits the expression of genes that respond to IL-4 and IL-13 (il4 and il13 activate stat6) by suppressing STAT6 activation.   1. Schülke S. Induction of Interleukin-10 Producing Dendritic Cells As a Tool to Suppress Allergen-Specific T Helper 2 Responses. Front Immunol. 2018 Mar 19;9:455. doi: 10.3389/fimmu.2018.00455. PMID: 29616018; PMCID: PMC5867300. | |  |
| NFkB | 1 | - IL1R & !Bcl3 |
| Activates the transcription of genes that code for cytokines such as IL-1B and TNFa and is involved in the production of other inflammatory cytokines such as interferon gamma (IFN-g), IL-6, IL-2, and IL-8.   1. López-Bojorquez, Lucia Nikolaia. (2004). La regulación del factor de transcripción NF-κB. Un mediador molecular en el proceso inflamatorio. Revista de investigación clínica, 56(1), 83-92. Recuperado en 22 de marzo de 2021, de http://www.scielo.org.mx/scielo.php?script=sci_arttext&pid=S0034-83762004000100012&lng=es&tlng=es. | |  |
| STAT6 | 1 | - !STAT3 |
| STAT6 is able to downregulate IL-10 production and promote IL-12 production.   1. <https://www.ncbi.nlm.nih.gov/pmc/articles/PMC3107597/> | |  |
| TCF7L2 | 1 | - TFCP2L1 & !CEBPB & !IRF8 |
| Related with Wnt signaling.  Our pattern matching results identified TFBS of TFCP2L1, CEBPB and IRF8   1. <https://link.springer.com/chapter/10.1007/5584_2021_663>   <https://pubmed.ncbi.nlm.nih.gov/30842629/> | |  |
| TFCP2L1 | 1 | - STAT3 |
| STAT3 activates TFCP2L1   1. <https://link.springer.com/chapter/10.1007/5584_2021_663>   <https://pubmed.ncbi.nlm.nih.gov/30842629/> | |  |
| IRF8 | 1 | - ERK |
| ERK signaling IRF8 activation   1. <https://pubmed.ncbi.nlm.nih.gov/34705053/> | |  |
| CEBPB | 1 | - NFkB |
| Active in inflamatory response.   1. <https://pubmed.ncbi.nlm.nih.gov/26526992/>   <https://pubmed.ncbi.nlm.nih.gov/30842629/> | |  |
| GAL4 | 1 | - TFCP2L1 & !CEBPB |
| Our pattern matching results identified TFBS of TFCP2L1 and CEBPB   1. <https://pubmed.ncbi.nlm.nih.gov/30842629/> | |  |
| SOCS3 | 1 | - STAT3 |
| SOCS-3 inhibits NFkB translocation into the nucleus and the associated induction of proinflammatory gene expression.  SOCS-3 also mediates Jak1 inhibition, resulting in feedback inhibition of the JAK1/Tyk2/STAT3 pathway  Other gene products that respond to STAT3 gene expression inhibit TLR4 signaling at the level of IRAK and TRAF6   1. Shouval, DS, Ouahed, J., Biswas, A., Goettel, JA, Horwitz, BH, Klein, C., Muise, AM y Snapper, SB (2014). Señalización del receptor de interleucina 10:regulador maestro de la homeostasis de la mucosa intestinal en ratones y humanos. Avances en inmunología , 122 , 177–210. https://doi.org/10.1016/B978-0-12-800267-4.00005-5 | |  |
| NFkBp50p50 | 1 | - IL10 |
| IL-10 promotes nuclear translocation of NF-kB-p50 / p50, promotes IL-10 expression in association with Bcl3. NFKB is thought to be activated by an IL10 receptor signaling pathway.   1. Comi M, Amodio G, Gregori S. Interleukin-10-Producing DC-10 Is a Unique Tool to Promote Tolerance Via Antigen-Specific T Regulatory Type 1 Cells. Front Immunol. 2018 Apr 6;9:682. doi: 10.3389/fimmu.2018.00682. PMID: 29686676; PMCID: PMC5900789. | |  |
| IL1RN | 1 | - STAT3 |
| IL-1RN is a decoy protein that binds to the IL-1 receptor, blocking proinflammatory signaling normally initiated by IL-1β binding to this receptor   1. Schülke S. Induction of Interleukin-10 Producing Dendritic Cells As a Tool to Suppress Allergen-Specific T Helper 2 Responses. Front Immunol. 2018 Mar 19;9:455. doi: 10.3389/fimmu.2018.00455. PMID: 29616018; PMCID: PMC5867300. | |  |
| IL1R | 1 | - !IL1RN |
| Once IL-1 binds to IL-1R, it triggers a cascade of kinases that produce a strong proinflammatory signal that leads to NFκB activation..   1. <https://www.ncbi.nlm.nih.gov/pmc/articles/PMC5756628/> | |  |
| Bcl3 | 1 | - STAT3 |
| Together with NFkBp50p50 it will promote the expression of IL-10. It has been seen that it inhibits the expression of TNF-a by inhibiting NFkBNFkB   1. Comi M, Amodio G, Gregori S. Interleukin-10-Producing DC-10 Is a Unique Tool to Promote Tolerance Via Antigen-Specific T Regulatory Type 1 Cells. Front Immunol. 2018 Apr 6;9:682. doi: 10.3389/fimmu.2018.00682. PMID: 29686676; PMCID: PMC5900789. | |  |
| INFsig | 1 | - NFkB |
| The production of proinflammatory cytokines (IL-1b, IL-6, tumor necrosis factor alpha) and the expression of both MHC II and costimulatory molecules (CD80, CD83, CD86) are increased.   1. Schülke S. Induction of Interleukin-10 Producing Dendritic Cells As a Tool to Suppress Allergen-Specific T Helper 2 Responses. Front Immunol. 2018 Mar 19;9:455. doi: 10.3389/fimmu.2018.00455. PMID: 29616018; PMCID: PMC5867300.   <https://www.ncbi.nlm.nih.gov/pmc/articles/PMC4741283/> | |  |
| IL12 | 1 | - STAT6 & !TFCP2L1 & !GAL4 & !TCF7L2 |
| PRO INFLAMATORY CYTOKINE   1. <https://pubmed.ncbi.nlm.nih.gov/30664357/>   <https://pubmed.ncbi.nlm.nih.gov/30842629/> | |  |
| IL6 | 1 | - CEBPB & !TFCP2L1 & !TCF7L2 & !GAL4 |
| PRO INFLAMATORY CYTOKINES   1. <https://doi.org/10.1016/j.cyto.2014.12.027>   <https://pubmed.ncbi.nlm.nih.gov/30842629/> | |  |
| TNF | 1 | - CEBPB & IRF8 & !TFCP2L1 |
| PRO INFLAMATORY CYTOKINES   1. <https://pubmed.ncbi.nlm.nih.gov/29298134/>   <https://pubmed.ncbi.nlm.nih.gov/30842629/> | |  |
| IL10_gene | 1 | - NFkBp50p50 & Bcl3 & !STAT6 |
| Inhibits TH1 cell differentiation by reducing IL2, IL12, and INFg, limits effector T cell function by suppressing TNFa, IL1B, and IL6.   1. <https://www.frontiersin.org/articles/10.3389/fimmu.2018.00455/full> | |  |
| C1QA | 1 | - TFCP2L1 & GAL4 & TCF7L2 |
| ANTI-INFLAMATORY   1. <https://pubmed.ncbi.nlm.nih.gov/18504288/>   <https://pubmed.ncbi.nlm.nih.gov/30842629/> | |  |
| THBS1 | 1 | - CEBPB & !TFCP2L1 & !GAL4 |
| ANTI-INFLAMATORY   1. <https://pubmed.ncbi.nlm.nih.gov/32517571/>   <https://pubmed.ncbi.nlm.nih.gov/30842629/> | |  |
| CCR5 | 1 | - TFCP2L1 & !CEBPB |
| CHEMOKINES   1. <https://pubmed.ncbi.nlm.nih.gov/19811270/>   <https://pubmed.ncbi.nlm.nih.gov/30842629/> | |  |
| CCR7 | 1 | - TFCP2L1 & !CEBPB |
| CHEMOKINES   1. <https://pubmed.ncbi.nlm.nih.gov/26858545/>   <https://pubmed.ncbi.nlm.nih.gov/30842629/> | |  |
| CXCR4 | 1 | - GAL4 |
| CHEMOKINES   1. <https://www.ncbi.nlm.nih.gov/pmc/articles/PMC2946082/>   <https://pubmed.ncbi.nlm.nih.gov/30842629/> | |  |
| CCL22 | 1 | - !CEBPB & TFCP2L1 & TCF7L2 |
| CHEMOKINES   1. <https://pubmed.ncbi.nlm.nih.gov/30842629/> | |  |
| CXCL1 | 1 | - TFCP2L1 & TCF7L2 & !IRF8 |
| CHEMOKINES   1. <https://www.ncbi.nlm.nih.gov/pmc/articles/PMC8711184/>   <https://pubmed.ncbi.nlm.nih.gov/30842629/> | |  |
| ALDH2 | 1 | - TFCP2L1 & GAL4 |
| METABOLIC CONTROL   1. <https://pubmed.ncbi.nlm.nih.gov/30842629/> | |  |
| ICOSLG | 1 | - TFCP2L1 & GAL4 |
| IMMUNOMODULATORY MOLECULES   1. <https://pubmed.ncbi.nlm.nih.gov/15476242/>   <https://pubmed.ncbi.nlm.nih.gov/30842629/> | |  |
| CTLA4 | 1 | - TFCP2L1 & GAL4 & !IRF8 |
| IMMUNOMODULATORY MOLECULES   1. <https://pubmed.ncbi.nlm.nih.gov/19950182/>   <https://pubmed.ncbi.nlm.nih.gov/30842629/> | |  |
| IL1B | 1 | - CEBPB & !TFCP2L1 & !TCF7L2 & !GAL4 |
| PRO INFLAMATORY CYTOKINES   1. <https://pubmed.ncbi.nlm.nih.gov/31082500/>   <https://pubmed.ncbi.nlm.nih.gov/30842629/> | |  |
| CD80 | 1 | - CEBPB & !TCF7L2 |
| DC MATURATION   1. <https://pubmed.ncbi.nlm.nih.gov/10023862/>   <https://pubmed.ncbi.nlm.nih.gov/30842629/> | |  |
| CD86 | 1 | - CEBPB & !TCF7L2 |
| DC MATURATION   1. <https://pubmed.ncbi.nlm.nih.gov/10023862/>   <https://pubmed.ncbi.nlm.nih.gov/30842629/> | |  |
| HLA_DR | 1 | - CEBPB & !GAL4 & !TCF7L2 |
| 1. <https://pubmed.ncbi.nlm.nih.gov/30842629/> | |  |
| IDO1 | 1 | - TCF7L2 & !CEBPB |
| METABOLIC CONTROL   1. <https://pubmed.ncbi.nlm.nih.gov/30170030/>   <https://pubmed.ncbi.nlm.nih.gov/30842629/> | |  |
| HMOX1 | 1 | - GAL4 & !IRF8 |
| METABOLIC CONTROL   1. <https://www.ncbi.nlm.nih.gov/pmc/articles/PMC3401980/>   <https://pubmed.ncbi.nlm.nih.gov/30842629/> | |  |
| ARG1 | 1 | - TCF7L2 & !CEBPB |
| METABOLIC CONTROL   1. <https://www.ncbi.nlm.nih.gov/pmc/articles/PMC4895207/>   <https://pubmed.ncbi.nlm.nih.gov/30842629/> | |  |
| CCL18 | 1 | - !IRF8 |
| CHEMOKINES   1. <https://pubmed.ncbi.nlm.nih.gov/30842629/> | |  |
| BTLA | 1 | - TCF7L2 & !CEBPB |
| IMMUNOMODULATORY MOLECULES   1. <https://www.ncbi.nlm.nih.gov/pmc/articles/PMC6770819/>   <https://pubmed.ncbi.nlm.nih.gov/30842629/> | |  |
| PDCD1 | 1 | - TCF7L2 |
| IMMUNOMODULATORY MOLECULES   1. <https://pubmed.ncbi.nlm.nih.gov/24860805/>   <https://pubmed.ncbi.nlm.nih.gov/30842629/> | |  |
| ERK | 1 | - !IL10 & !SOCS3 |
| MAP kinases, also known as extracellular signal-regulated kinases (ERKs), act as an integration point for multiple biochemical signals, and are involved in a wide variety of cellular processes such as proliferation, differentiation, transcription regulation and development.   1. <https://www.genecards.org/cgi-bin/carddisp.pl?gene=MAPK1&keywords=erk1%2F2> | |  |
| FASLG | 1 | - !CEBPB |
| IMMUNOMODULATORY MOLECULES   1. <https://pubmed.ncbi.nlm.nih.gov/27050822/> | |  |
| CD40 | 1 | - GAL4 & TCF7L2 |
| DC MATURATION   1. <https://pubmed.ncbi.nlm.nih.gov/26289938/>   <https://pubmed.ncbi.nlm.nih.gov/30842629/> | |  |
| IRF4 | 1 | - !GAL4 & !TFCP2L1 & !TCF7L2 |
| Homogeneous differential gene expression across the datasets, identifying 53 genes with a combined p-value<10^5 which we deemed to be characteristic of tolDC. Downregulated Genes: IRF4, CD1A, CD1E, CD1B, CD1C.   1. <https://pubmed.ncbi.nlm.nih.gov/34745103/> | |  |
| CD1C | 1 | - !TFCP2L1 & !TCF7L2 |
| Homogeneous differential gene expression across the datasets, identifying 53 genes with a combined p-value<10^5 which we deemed to be characteristic of tolDC. Downregulated Genes: IRF4, CD1A, CD1E, CD1B, CD1C.   1. <https://pubmed.ncbi.nlm.nih.gov/34745103/> | |  |
| CD1E | 1 | - CEBPB & !TFCP2L1 |
| Homogeneous differential gene expression across the datasets, identifying 53 genes with a combined p-value<10^5 which we deemed to be characteristic of tolDC. Downregulated Genes: IRF4, CD1A, CD1E, CD1B, CD1C.   1. <https://pubmed.ncbi.nlm.nih.gov/34745103/> | |  |
| CD1B | 1 | - CEBPB & IRF8 & !GAL4 |
| Homogeneous differential gene expression across the datasets, identifying 53 genes with a combined p-value<10^5 which we deemed to be characteristic of tolDC. Downregulated Genes: IRF4, CD1A, CD1E, CD1B, CD1C.   1. <https://pubmed.ncbi.nlm.nih.gov/34745103/> | |  |
| CD1A | 1 | - !GAL4 & !TFCP2L1 |
| Homogeneous differential gene expression across the datasets, identifying 53 genes with a combined p-value<10^5 which we deemed to be characteristic of tolDC. Downregulated Genes: IRF4, CD1A, CD1E, CD1B, CD1C.   1. <https://pubmed.ncbi.nlm.nih.gov/34745103/> | |  |
